# Exploring the interaction between the pupillary light response after exogenous attentional cueing and detection performance

**DOI:** 10.1101/2020.04.08.032060

**Authors:** Tiemen J. Wagenvoort, Rosanne H. Timmerman, Stefan Van der Stigchel, Jasper H. Fabius

## Abstract

Pupil size changes under different light conditions. Whereas this pupillary light response (PLR) has long been regarded to be influenced by luminance only, recent studies indicated the PLR is also modulated by cognitive factors such as the allocation of spatial attention. This attentional modulation of the PLR has previously been hypothesized to facilitate detection and discrimination of visual information. Here, we replicated the finding that the pupil dilates when a cue is presented at the dark side of a screen and constricts when the cue is presented at the bright side, even when the eyes are fixated at the center. Furthermore, we investigated whether this modulation of the PLR, evoked by exogenous shifts of covert attention, facilitates perception operationalized as detection performance for threshold stimuli. Results showed that a larger pupil was indeed related to increased detection performance, although this effect was restricted to conditions in which both cue and target appeared on a dark surface. Our findings are in line with the notion that pupil dilations improve detectability, whereas pupil constrictions enhance discriminability of small stimuli.

## Introduction

The pupillary light response (PLR) is a basic reflex of the human body. The pupil dilates (i.e. becomes larger) when the eyes are fixating in a dark environment and constricts (i.e. becomes smaller) for bright environments (Beatty & Lucero-Wagoner, 2000). Simply put, a dilated pupil causes more light to enter the eye and subsequently allows for better perception in the dark (Woodhouse & Campbell, 1975). On the other hand, a constricted pupil increases depth of field (Campbell, 1957) and visual acuity (Campbell & Gregory, 1960; Woodhouse, 1975).

The PLR was long regarded to be influenced by luminance only (Loewenfeld, 1958). However, recent studies indicate that it can be influenced by cognitive factors as well (Hartmann & Fischer, 2014; Mathôt & Van der Stigchel, 2015). For example, besides luminance, perceived brightness also influences pupil size (Binda, Pereverzeva, & Murray, 2013b; Laeng & Endestad, 2012; Naber & Nakayama, 2013): the pupil was shown to constrict more when subjects looked at an image of a lamp compared to an image of a rock, even though both images had the same luminance. Furthermore, cognitive processes such as emotion (Hess & Polt, 1960), mental load (Hess & Polt, 1964), arousal (Bradshaw, 1967) and memory (Kahneman & Beatty, 1966) were found to be related to the PLR. Recently, it has even been demonstrated that pupil size is indicative of the location of covert attention i.e. shifting attention without making eye movements (Binda & Murray, 2015; Binda, Pereverzeva, & Murray, 2013a; Canty, 2002; Mathôt, Dalmaijer, Grainger, & Van der Stigchel, 2014; Mathôt, van der Linden, Grainger, & Vitu, 2013). For instance, when participants fixated on the center of a screen with a bright and dark part, their pupils relatively constricted when they covertly attended to the bright part, and relatively dilated when they covertly attended on the dark part (Binda et al., 2013a; Mathôt et al., 2014, 2013). This phenomenon can be elicited both by endogenous (Binda et al., 2013a; Mathôt et al., 2013) and exogenous cueing (Fabius, Mathôt, Schut, Nijboer, & Van der Stigchel, 2017; Mathôt et al., 2014). Moreover, this effect appeared to reverse after approximately one second: when initially on one of the sides, covert attention shifted to the other side after a period of time, reminiscent of inhibition of return (IOR).

It is currently unknown whether the PLR actually serves a perceptual function (Mathôt, 2018), but it is tempting to think that it does. A typical hypothesis is that the PLR, evoked by the aforementioned effects of cognitive processes, also benefits perception (Binda & Murray, 2015; Mathôt & Van der Stigchel, 2015; Wang & Munoz, 2015). Meanwhile, this speculation is nuanced by considering that pupillary reactions are found to be relatively slow (Beatty & Lucero-Wagoner, 2000) compared to the appearance of certain stimuli. The average time to peak dilation of ∼0.9 seconds (Hoeks & Levelt, 1993) might not be sufficient to facilitate for perception of stimuli that appear in the order of milliseconds.

In the present study, we investigated whether the PLR, evoked by exogenous shifts of covert attention, facilitates detection performance for threshold stimuli. To measure the effect of covert attention on the pupil, we examined the difference in pupil size between cues on a dark background minus pupil size between cues on a bright background. To measure the behavioral effect of the cue on perception, we constructed a detection performance time series using a dense sampling, exogenous cueing paradigm (Landau & Fries, 2012). If covert attention influences the PLR, the pupil should relatively dilate if a cue is presented at the dark side of a screen, and constrict if the cue is presented at the bright side of a screen, similar to previous studies (Fabius et al., 2017; Mathôt et al., 2014). Furthermore, we hypothesized that if targets appear at the same side of the screen as the preceding cue early in the time series, these targets would be more likely to be detected compared to targets later in the time series or targets appearing at the other side of the screen (Mulckhuyse & Theeuwes, 2010). Lastly, if the PLR facilitates perception, detection of stimuli should be enhanced at the same time at which the pupil size is affected by the background color of the cue.

## Methods

### Participants

There were 20 participants (11 female, mean age 23.8, range = [21, 43]). All participants reported normal or corrected-to-normal vision. All experimental procedures were approved by the local ethical committee of the Faculty of Social Science of Utrecht University.

### Materials

The experiment took place in a dark room. Participants were placed in a chinrest to ensure stability of the head. Responses were given by pressing keys on the keyboard. Two fingers of the dominant hand of the participants were placed on the keys ‘V’ and ‘N’. An Asus RoG Swift PG278Q LCD display was used for the experiment. Screen contrast was set on 50/100 and screen brightness on 0/100. ULMB was turned off and the frequency of the screen was set on 100 Hz. We did not linearize the screen, because that disabled us from showing targets with low luminance on the dark side of the display. An Eyelink 1000 (SR Research, Ltd., Ottawa, ON) was used as eye tracker, which sampled at 1000 Hz. Only the left eye was tracked. The experiment was programmed in Matlab (R2017a, The Math Works, Inc., Natick, MA), with the Psychtoolbox (Brainard, 1997) and the Eyelink toolbox (Cornelissen, Peters, & Palmer, 2002).

### Stimuli

During the experiment, the screen was divided into three segments (Figure 1). The left part of the screen was bright (53 cd/m^2^). A grey placeholder (13.8 cd/m^2^) of 10° of visual angle in diameter was positioned on the bright part 8° of visual angle to the left of the center of the screen. The right part of the screen was dark (0.15 cd/m^2^). Another grey ring, serving as a placeholder, was positioned on the dark part 8° of visual angle to the right of the center of the screen. The placeholders indicated in which regions a target could appear. The bright and dark parts were separated by a grey bar (13.8 cd/m^2^) with a width of 5.4° visual angle. The purpose of this grey bar was to minimize the change in illumination when participants made microsaccades and other minor eye movements. On top of the grey bar, a green (18.3 cd/m^2^) fixation cross superimposed on a fixation dot (diameter = 0.5°) was presented in the center of the screen (Thaler, Schütz, Goodale, & Gegenfurtner, 2013). We flipped the screen layout randomly between participants, which resulted in 9 participants having the bright side on the left, and 11 participants having the bright side on the right side of the screen.

**Figure 1.**
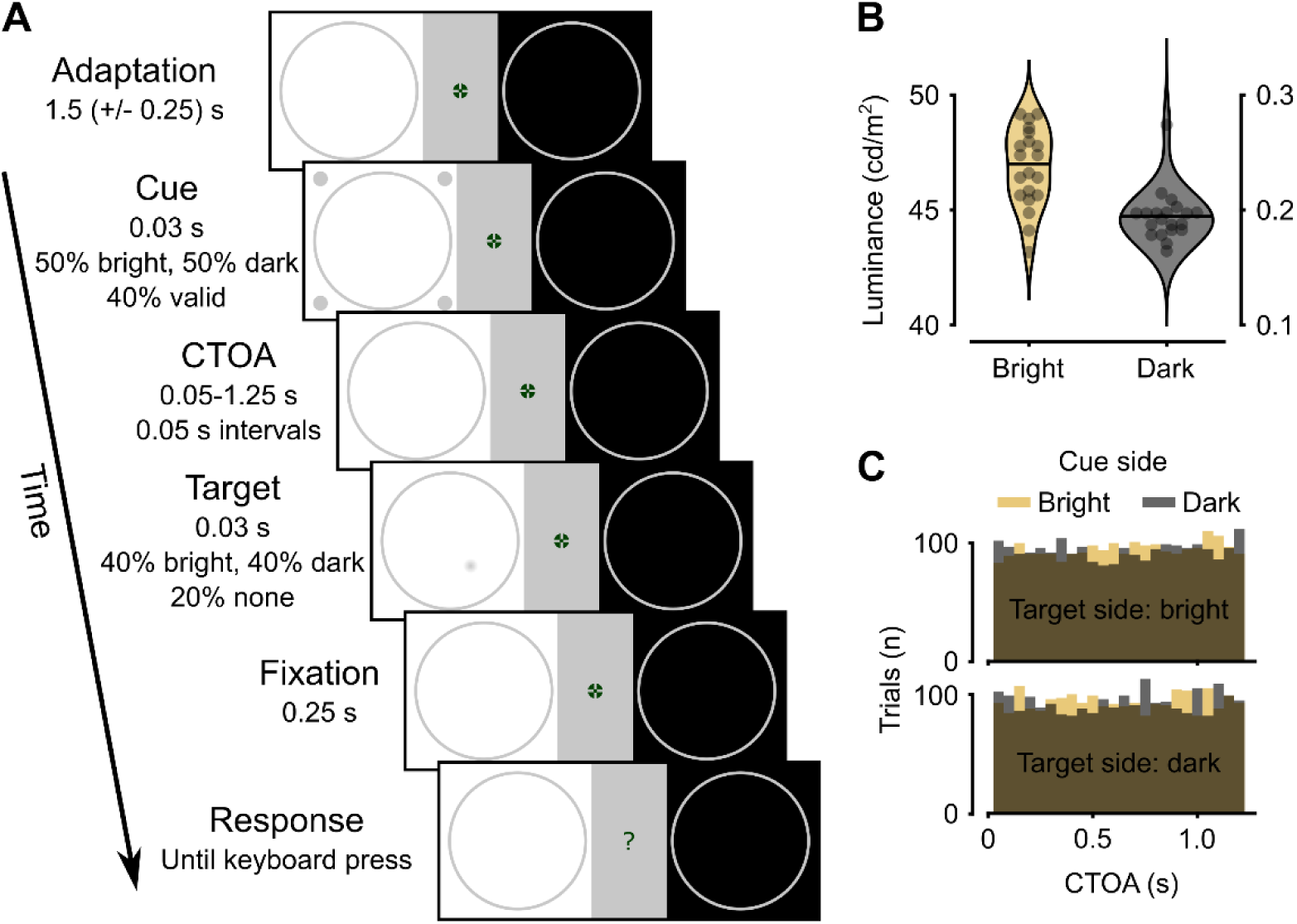
**A**. Visualization of a single trial. Two grey circles (placeholders) were presented on a partly bright (53 cd/m2), partly dark screen (0.15 cd/m2), separated by a grey bar (13.8 cd/m2), i.e. the adaptation screen (Mathôt et al., 2014). We flipped the dark and bright parts randomly between participants. It was presented for 1.5 (+/- 0.25) s before cue onset. The cue consisted of four dots appearing around one of the two circles for 30 ms, with 40% validity. The cue was followed by a cue-target-onset-asynchrony (CTOA) between 0 and 1.25 s, in 50 ms intervals. In 80% of the trials, a target was presented in the left or the right circle. In 20% of the trials there was no target. The fixation screen was presented again for 0.25 ms to prevent masking. In the response screen the fixation cross changed into a question mark. Participants indicated whether a target was present or not. **B**. Luminance thresholds estimates from the staircase session before the experiment. The procedure of the trials in the staircase session was the same as displayed in A, with the exception that there was no cue. Luminance was adjusted with QUEST. We used two staircases of 32 trials for each side of the display. Target luminance in the experiment was set as the average of the estimates of these two staircases. Each dot per violin represents the target luminance of one subject. **C**. Number of trials per CTOA and per condition across all 20 participants after trial exclusion. CTOA’s were uniformly distributed across bins.

Attention was attracted to one side of the screen using exogenous cueing (Posner, 1980). The cues consisted of four grey dots (0.5° visual angle in diameter, 13.8 cd/m^2^) and were placed around the placeholders at 1.5° visual angle from the edge of the placeholders (Landau & Fries, 2012). Targets were small (s.d. = 0.25°) luminance increments when presented on a dark background or decrements when presented on a bright background that appeared at a random location within one of the placeholders. The amplitude of the luminance modulation was fixed at the 70% detection threshold. These thresholds were measured before the start of the experiment using two interleaved QUEST procedures (Watson & Pelli, 1983) per background luminance, i.e. four in total (see *Procedure*).

### Procedure

Before starting the experiment, participants provided informed consent. Then, a nine-point calibration of the eye tracker was performed. During this calibration, participants fixated on a dot until it reappeared at another location. Validation of this calibration had to be rated as ‘good’ to proceed. The calibration and validation procedure were repeated when deemed necessary by the experimenter. The experiment consisted of three parts: a practice session (20 trials over 1 block), a staircase procedure to assess target luminance at a performance of p_detected_ = 0.7 (160 trials over 8 blocks) and the actual experiment (640 trials over 32 blocks). Between blocks, participants could take a short rest when needed. In total, the experiment lasted approximately two hours.

### Experimental procedure

As shown in Figure 1, a single trial consisted of six parts. First, each trial began with an adaptation period of 1.5 seconds (+/- 0.25 s, uniformly distributed) to stabilize pupil size at a baseline level before cue onset (Mathôt et al., 2014). Second, the cue was presented for 30 ms around one of the placeholders (similar to Landau & Fries, 2012). Third, target onset was delayed with respect to cue offset with a delay between 0 and 1.25 seconds, with intervals of 0.05 s. Because of the 100 Hz refresh rate of the screen, a target could be shown at one of the 25 equally spaced intervals in the 1.25 s target period of a trial. All 25 intervals were used five times in order to get a detailed view of detection performance over time. Fourth, the target appeared. In 80% of the trials, the target was presented for 30 ms. The remaining 20% were target-absent ‘catch’ trials, used to estimate false alarm rates (Busch, Dubois, & VanRullen, 2009). Because targets appeared on 80% of all trials and cue- and target-side were uncoupled, cue validity was (0.5×80% =) 40%. Additionally, the presence of 20% catch trials, combined with the thresholded target luminance (see *Staircase procedure*), meant that we expected participants to detect targets on (0.7×80%=) 56% of all trials. Fifth, a fixation screen was presented for 0.25 s. This screen made sure no target would appear just before the response phase (indicated by the fixation cross changing into a question mark), which could lead to masking effects (Marcel, 1983). Sixth, the fixation cross changed into a question mark, indicating that a response (target detected yes/no) had to be given. This screen lasted until a keyboard response was given. After a response was provided, the next trial started immediately. During the experiment, eye position was checked online. When the eye drifted away for more than 2.5**°** visual angles from the fixation cross during the target period, the central fixation cross changed into an X. Also, an X appeared when participants blinked during the target presentation period. In these cases, the trial was repeated.

### Staircase procedure

A QUEST routine (Watson & Pelli, 1983) was used prior to the experiment to adjust the luminance levels of the detection targets per participant to have all participants perform around a detection performance of p_detected_ = 0.7. Four interleaved QUEST staircases were used (two per background luminance). Each staircase was completed using 160 trials. Detection probability was modelled as a function of the base 10 logarithm of Weber contrast (between 0 and 1) with a Weibull link function. The parameters for this hypothetical underlying psychometric function were: β = 3.5, δ = 0.03, γ = 0 for all participants. The initial threshold guess was set at a Weber contrast of 0.08. The final luminance level of the stimulus was taken as the average thresholds estimate of the two staircases per background color. In addition to the trials with targets, we also included catch trials (20% of total amount of trials), in which no target was displayed during the staircase procedure. These trials were not used to update the threshold estimate, rather we used the catch trials to assess the false alarm rate. In this staircase procedure the same drifting and blinking criteria as in the experimental procedure were applied.

### Practice

To ensure that the participants understood the experiment, there was a practice session before the staircase procedure. Participants could practice for one block. Target detection in the practice session was relatively easy (i.e. Weber contrast of 0.3) as compared to the rest of the experiment, since there had not been a staircase procedure yet. Consequently, the intensity of the targets was very high. Apart from the intensity of the targets, the practice session was the same as the actual experiment.

## Analysis

### Preprocessing

We only included trials that contained no saccades with amplitudes > 1° between cue onset and the onset of the response window. Saccades were detected using the algorithm of Nyström & Holmqvist (2010). Moreover, we did not include trials with missing samples in the pupil time series between cue onset and the response window. Furthermore, we only included trials in which responses were given between 0.2 and 2.0 s after the fixation cross changed into a question mark. After saccade detection, eye-movement data and pupil size was downsampled to 100 Hz. We extracted epochs from the pupil time series from -1.25 ms before, to 2.06 ms after cue onset. All pupil sizes were normalized by subtracting the average pupil size in the window of -0.2 to 0 ms before cue onset (Mathôt, Fabius, Heusden, & Stigchel, 2018).

### Statistics

We performed three analyses: (1) on the pupil size as a function of cue location (dark/bright), (2) on the behavioral cueing effect as a function of congruency, and (3) on the relationship between pupil size and detection performance.

To investigate (1) pupil size as a function of cue location, we used linear mixed effects models (Bates, Mächler, Bolker, & Walker, 2015). At each time point, we modelled normalized pupil size as a function of cue-side (dark vs. bright) with random intercepts and slopes per participant. In Wilkinson’s notation: *pupil size ∼ cue side + (1+cue side* | *participant)*. Each time point of the pupil size time series was analyzed with this model. To increase the number of trials per time point, we also included trials from the two time points before and two time points after the current, i.e. a temporal smoothing of 2 samples or 0.1 s. To correct for multiple comparisons, we used threshold-free cluster-enhancement (TFCE) and 5000 permutation tests (Maris & Oostenveld, 2007; Smith & Nichols, 2009). Briefly, this entails running the same statistical models but with the labels of cue side permuted. The permutations were stratified over participants, i.e. each permuted data set contained the same overall responses of each subject and the same number of trials per condition per participant. For each permutation, we summed F-values of clusters where the F-values were larger than the critical F-value (with alpha = 0.05); the critical F-value differs slightly between different cue target onset asynchronies (CTOA’s) because the number of trials differs slightly between CTOA’s. The permutation method gives a distribution of F-values based on the permuted datasets. We then determined whether the summed F-values of clusters in the original dataset were larger than the 95^th^ percentile of the distribution of F-values from the permutations.

To investigate (2) the behavioral cueing effect as a function of congruency, we used linear mixed effects models with a logistic link function. Responses were defined as hits and misses (i.e. we only used trials where a target was presented). Responses were modelled as a function of side (bright vs. dark) and cue-target congruency (congruent vs. incongruent) and their interaction. In Wilkinson’s notation: *response ∼ cue side * congruency + (1 + cue side * congruency* | *participant)*. Similar to the analysis of pupil size, we applied a temporal smoothing window of 0.1 s and corrected for multiple comparisons using 5000 permutations and TFCE.

We investigated (3) the relation between pupil size and detection performance also using a logistic linear mixed effects model per time point. Specifically, we modelled detection performance (hit vs. miss) as a function of three main factors: cue side (bright vs. dark), target side (bright vs. dark) and pupil size (relative to baseline), with a logistic link function. In addition to these main effects, we also included their interactions. For the random effects, we added random intercepts and slopes for each main and interaction effect per participant. In Wilkinson’s notation: *detection ∼ cue side * target side * pupil size + (1 + cue side * target side * pupil size* | *participant)*. For this analysis, we also applied a temporal smoothing window of 0.1 s and corrected for multiple comparisons using 5000 permutations and TFCE.

In short, interpretation of this mixed effect is as follows. We set the baseline to be trials with a cue on the bright side, a target on the bright side and a pupil size of zero. So, the intercept of the model gives the detection performance for trials with both a cue and a target on the bright side of the screen and a normalized pupil size of zero. A main effect of cue side, for example, would mean that detection performance was higher after a cue on the dark side than on the bright side, when the target appeared on the bright side. Negative parameter estimates of this factor could be interpreted as a cueing effect: targets on the bright side would be less likely to be detected when pupil size was relatively large. For the three-way interaction effect (cue side, target side, pupil size) positive parameters would indicate higher detection performance when the cue and target appeared on the dark side whilst pupil size was relatively large.

## Results

### False alarms

As a general check for whether participants were performing our task as intended, we tested for differences in false alarm rates between trials with cues on the bright vs. trials with cues on the dark side with a paired t-test. False alarm rates were converted to log-odds. In case of 0 false alarms, the rate was adjusted with a log-linear approach (i.e. adding 0.5 and dividing by the number of trials without a target + 1). The median false alarm rate for trials with a cue on the bright side was 0.026 (range = [0.008, 0.246]). The median false alarm rate for trials with a cue on the dark side was 0.016 (range = [0.008, 0.349]). There was no significant difference between the two conditions (t(19) = 0.702, p = 0.49).

### Pupillary cueing effect

We examined the effect of the cue on pupil size, without considering subsequent behavioral performance. Overall, the pupil dilated after cue onset (Figure 2A). From 0.50 to 0.96 s after cue onset, the pupil dilated more in the cue on dark condition compared to the cue on bright condition (Figure 2B, maximum average relative dilation = 40.9 a.u., p_cluster_ < 0.0001). This relative increase seemed to reverse around 1.1 s (Figure 2B), but no statistical significance was reached.

**Figure 2.**
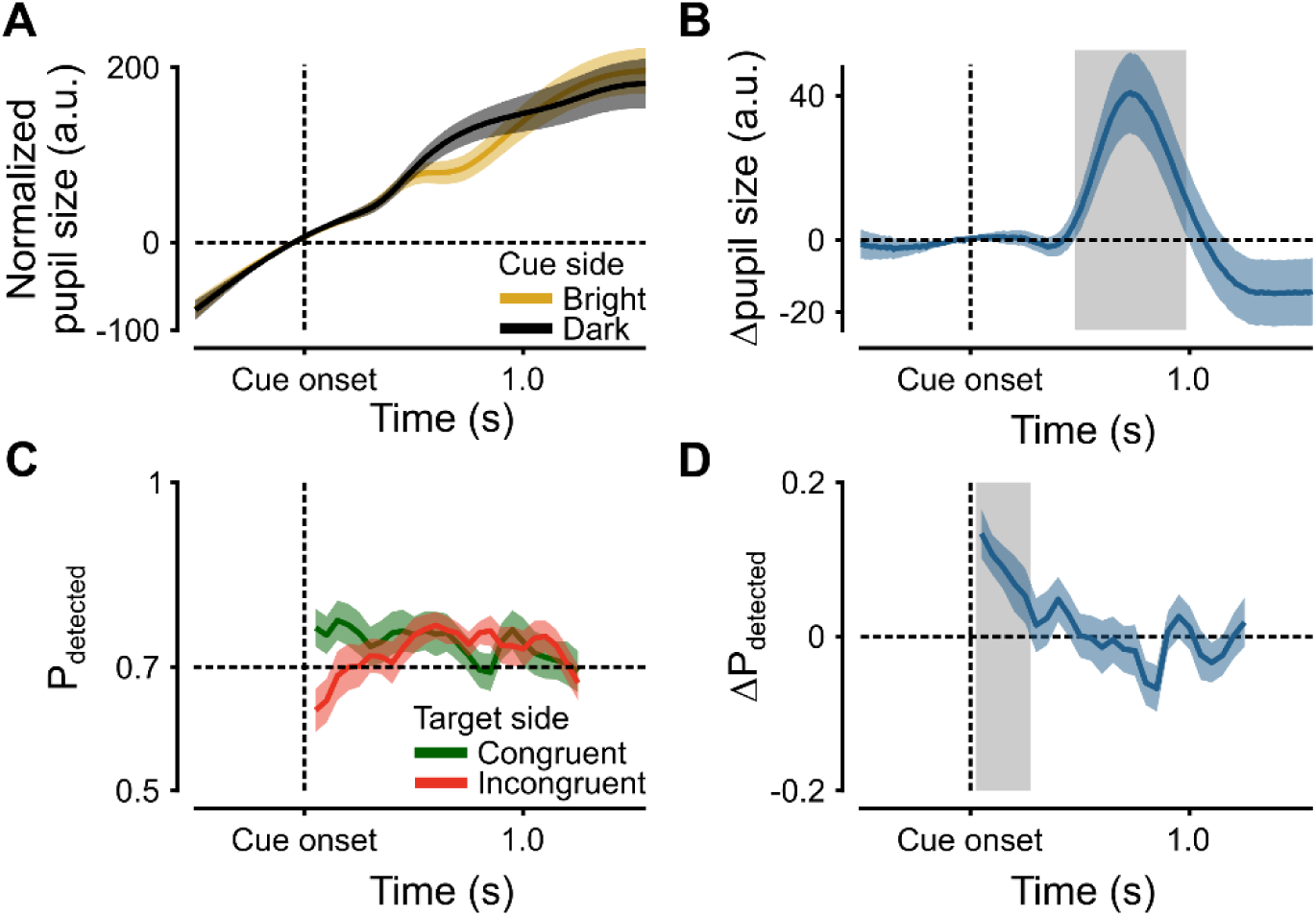
Main results. **A**. Normalized pupil size (arbitrary units) as a function of time after cue onset (seconds) for trials in which the cue appeared at the bright (yellow) and dark (black) part of the screen. Shaded areas indicate standard error of the mean across participants. **B**. Relative pupil size (dark - bright). Grey patch (t = 0.50 - 0.96) indicates a period with a significant difference between pupil size dark and bright. **C**. Detection performance (proportion detected) for all 25 CTOA’s for congruent (green) and incongruent (red) trials. **D**. Cueing effect for detection performance (difference in proportion detected). Grey patch (t = 0.05 - 0.25) show a significant difference between congruent and incongruent trials.

### Behavioral cueing effect

Next, we analyzed detection performance as a function of the congruency between the cue side and target side, for each CTOA. Detection performance differed for congruent trials compared to incongruent trials (Figure 2C). A significant positive cueing effect was found in detection performance for CTOA’s from 0.05 to 0.25 s (Figure 2D), indicating that detection performance was higher for congruent cue-target pairs compared to incongruent pairs (maximum group average ΔP_detected_ = 0.13, p_cluster_ < 0.0001). No later effects were observed.

### Interaction between pupil size and detection

We analyzed the interaction between pupil size and detection performance using a logistic linear mixed effects model. Please note that the unit of the reported parameter estimate (β) of the intercept is in log odds, whereas the units of all other parameter estimates are deviations from the intercept in log odds.

First, the effects of cue side and target side in the mixed effects model provide a re-analysis of the behavioral cueing paragraph above. This is a re-analysis because they describe detection performance as a function of cue side and target side (and their interaction) similar to the analysis described in the previous paragraph. The intercept was significantly above chance for all except the last CTOA (Figure 3A), indicating that targets on the bright side were reliably detected above chance after a cue on the bright side (mean β = 0.91, p_cluster_ = 0.0002). This shows that target intensity as determined by the staircase procedure was successfully set above the detection threshold of the participants.

**Figure 3.**
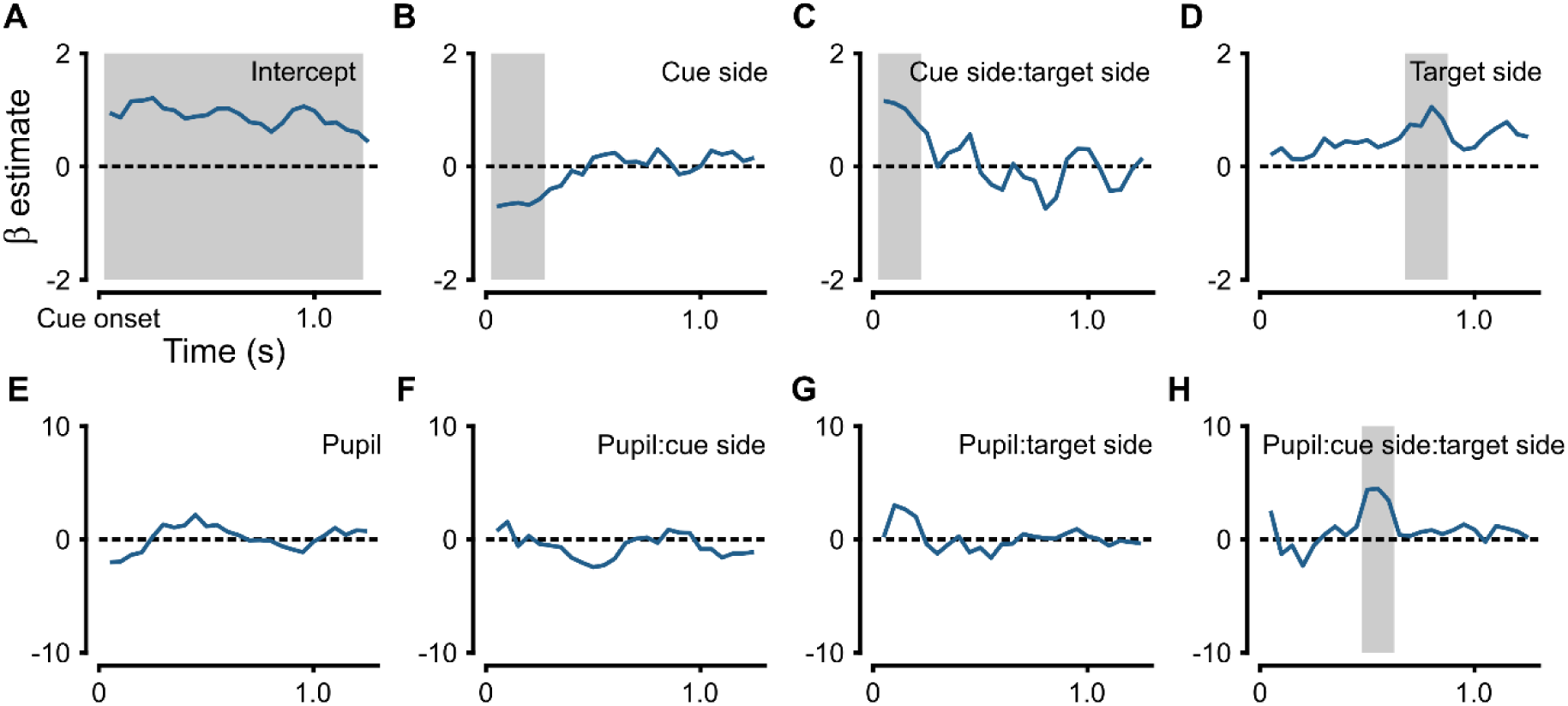
Result of logistic linear mixed effects model, modelling detection as a function of cue side (bright/dark), target side (bright/dark) and pupil size (a.u.). Models were made for each time point with a moving window of 3 time points (i.e. including the pre- and succeeding time points). Parameter estimates (β) are in log-odds relative to the intercept. Significance (grey patches) of the parameter estimates were corrected for multiple comparisons using threshold-free cluster-based permutation testing (N = 5000). **A**. Intercept i.e. log odds of detection in trials where both the target and cue were presented on the bright side of the screen. **B**. Detection performance was significantly lower than the intercept from cue onset to 0.25 s when the cue was presented on the dark side and the target on the bright side. **C**. Detection performance was significantly higher than the intercept until 0.20 s after cue onset when both the target and cue were presented on the dark side. **D**. Detection performance was significantly higher than the intercept from 0.70 to 0.85 s after cue onset when the cue was on the bright side and targets on the dark side. **E**. There was no significant modulation of pupil size on detection performance when the cue was presented on the bright side and the target on the bright side, (**F**.) nor when the cue was presented on the dark side and the target on the bright side, (**G**.) nor when the cue was presented on the bright side and the target on the dark side. **H**. Detection performance was significantly increased with increased pupil size in trials where both the cue and target were presented on the dark side between 0.50 - 0.60 s. after cue onset.

There was a negative modulation of detection performance by cue side (Figure 3B), with a significant modulation between cue onset to 0.25 s (mean β = -0.65, p_cluster_ < 0.0001). So, in that period, targets on the bright side were less likely to be detected when the cue had appeared on the dark side compared to when the cue had appeared on the bright side. Similarly, there was a positive modulation of detection performance between cue onset and 0.20 s (mean β = 1.02, p_cluster_ < 0.0001) when both the target and cue appeared on the dark side (Figure 3C), indicating that targets on the dark side were more likely to be detected when the cue had also appeared on the dark side. The time course of these effects, as displayed in Figure 3B and 3C describe the same positive cueing effect in detection performance as displayed in Figure 2C and 2D, albeit analyzed differently. In addition to this positive cueing effect, there was also a positive modulation on detection performance by target side between 0.70 and 0.85 s after cue onset (Figure 3D, mean β = 0.84, p_cluster_ < 0.0001). This positive modulation indicates that in that period after a cue on the bright side, targets on the dark side (incongruent) were significantly more likely to be detected than targets on the bright side (congruent). This effect resembles inhibition of return (Posner & Cohen, 1984; Posner, Rafal, Choate, & Vaughan, 1985). However, the demonstration of IOR seems to be limited to the bright side; it did not reach the significance level of alpha = 0.05 for targets and cues on the dark side (Figure 3B and 3C), neither did it appear in the analysis displayed in Figure 2C and 2D.

Next, we analyzed modulations of pupil size on detection performance. For targets on the bright side that appeared after a cue on the bright side, there was no significant modulation on detection performance by pupil size (Figure 3E). Neither did pupil size modulate detection performance when the cue was on the dark side and the target on the bright side (Figure 3F), nor when the cue was on the bright side and the target on dark side (Figure 3G). However, for targets that appeared on the dark side 0.50 to 0.60 s after a cue on the dark side (Figure 3H), detection performance increased with increasing pupil size (mean β = 4.11, p_cluster_ = 0.0018). This interval coincides with the beginning of the interval in which we observed a significant modulation of pupil size by cueing side (which is from 0.50 to 0.96 s, Figure 2B).

### Horizontal gaze position

To control for the effects of eye position, we also analyzed the horizontal gaze position with the same analysis as presented under *Pupillary cueing effect*. The average difference in horizontal gaze position between the two cue sides across participants never exceed 0.2° in either direction. At no point in the time series was there a significant effect of cue location on the horizontal gaze position.

## Discussion

It is generally assumed that the effect of attention on the PLR has a facilitating function (Binda & Murray, 2015; Mathôt, 2018; Mathôt & Van der Stigchel, 2015; Wang & Munoz, 2015). In this study, we investigated whether the PLR, evoked by exogenous attentional cues, facilitates detection of threshold stimuli. We expected that if covert attention influences the PLR, the pupil should dilate more if the cue was on the dark side of a screen than when the cue was on the bright side (i.e. the pupillary cueing effect). Furthermore, we hypothesized that if targets appeared at the same side of the screen as the cue, they would be more likely to be detected when the CTOA is short, compared to longer CTOAs or trials where cue and target appeared on different sides of the screen (i.e. behavioral cueing effect). Lastly, we examined whether the pupillary cueing effect modulated detection performance by testing whether detection performance increased if the pupil dilated and a target appeared at the dark side, and vice versa.

### Pupillary cueing effect

In general, the pupil dilated before and after cue onset (i.e. the positive trend visible in Figure 2A), which can be explained by an increase in arousal (Mathôt et al., 2014) and an orienting response (Wang, Boehnke, White, & Munoz, 2012). The pupillary cueing effect was apparent 0.50 s after cue presentation: pupil size differed between the cue on bright and cue on dark conditions (Binda & Murray, 2015; Mathôt et al., 2014, 2013). The pupil was relatively dilated when covert attention was exogenously cued to the dark part of the screen, and relatively constricted when cued to the bright part. Although the pupillary cueing effect seems to reverse at 1.10 s after cue onset, it does not reach significance. Hence, the data here do not provide evidence for pupillary IOR (as observed in Mathôt et al., 2014).

Based on this finding and supported by similar results of our pilot study (see Appendix), we conclude that our study further substantiates the notion that the PLR responds not just to luminance, but also has an attentional component (Mathôt & Van der Stigchel, 2015). More specifically, it highlights the relationship between pupil size and covert attention, by showing that the pupil is relatively dilated when covert attention is allocated on a dark surface, and relatively constricted when attention is allocated on a bright surface. It is important to mention that this effect occurs even though there was no actual change in brightness between both conditions, and no eye movements with amplitudes > 1° were made.

### Behavioral cueing effect

Immediately after cue onset, detection performance was higher in the congruent trials compared to the incongruent trials, indicating an increased likelihood of targets being detected when they followed a cue at the same side compared to the other side. This cueing effect can be explained by covert attention being attracted by the exogenous cue to a specific side, allowing for better detection of a subsequent target at the same side (congruent). On the contrary, a target appearing at the other side of a cue (incongruent) was more likely to be missed. This behavioral cueing effect did not persist after 0.25 s after cue onset. We found subtle IOR in detection performance (Posner & Cohen, 1984; Posner et al., 1985), although IOR was only apparent when a cue was given on the bright side of the screen, and only for a brief period.

### Modulation of detection performance by the pupillary cueing effect

To investigate the ability of the pupillary cueing effect to facilitate perception, we analyzed the relation between pupil size and detection performance. We found no relation between detection performance and pupil size when either the cue (Figure 3E), the target (Figure 3F) or both (Figure 3G) were presented on the bright side of the screen. Only when both the target and cue were presented on the dark side of the screen, we observed a significant modulation of pupil size on detection performance between 0.50 to 0.60 s after cue onset (Figure 3H). Around this time larger pupil sizes were related to higher detection performance. Interestingly, this period coincided with the beginning of the pupillary cueing effect as depicted in Figure 2B. Together, these findings suggest that the pupillary cueing effect facilitates detection performance on a dark background.

One possible explanation for the specificity of the modulation of the pupillary cueing effect could be that that, in general, pupil dilations result in enhanced sensitivity for faint stimuli in darkness (Aston-Jones & Cohen, 2005; Mathôt & Ivanov, 2019; Mathôt & Van der Stigchel, 2015). When in the current experiment covert attention was cued to a bright part, the pupil dilated compared to its state before cue onset. But when covert attention was cued to a dark part, the pupil dilated even more compared to before cue onset and compared to when attention was cued to a bright part. This additional increase in pupil size caused more light to enter the eye (Woodhouse & Campbell, 1975) and can have significantly affected the ability to detect threshold stimuli when they appeared at the dark part.

The enhanced detection performance for larger pupils resonates well with previous studies on pupil size and detection performance. For example, Mathôt and Ivanov (2019) had participants perform subsequent detection (in the central visual field) and discrimination (in the peripheral visual field) tasks. During these tasks they manipulated the overall background luminance across different blocks. Their data showed that detection performance is enhanced when pupil size is large and, conversely, discrimination performance is enhanced when pupil size is small.

With this study we further substantiate the notion that the PLR changes not only to overall luminance levels, but also to cognitive factors such as spatial attention. More specifically, it highlights the relationship between pupil size and covert attention by showing that the pupil dilates if a cue is presented at a dark part and constricts if a cue is presented at a bright part. We also provided evidence for the idea that a dilated pupil, caused by exogenous cues at a dark surface, enhances the likelihood of threshold targets being detected at this dark surface. We did not find support for the same phenomenon on the bright side. Possibly, this is because larger pupil sizes facilitate the detection of faint stimuli in dark environments, but small pupils do not in bright environments.

## Data availability

All data and experiment/analysis scripts for MATLAB are publicly available on Open Science Framework: https://osf.io/485mw/?view_only=b9d6eafde2b3459b833f540ed23056c3.

## Acknowledgements

This work was supported by The Netherlands Organization for Scientific Research VIDI Grant 452- 13-008 to SvdS. The authors declare no competing financial interests.

## Author contributions

**TJW**: Conceptualization, Methodology, Validation, Data curation, Writing - original draft, Project administration. **RHT**: Conceptualization, Methodology, Validation, Data curation, Writing - review & editing, Project administration. **SvdS**: Conceptualization, Writing - review & editing, Supervision. **JHF**: Conceptualization, Methodology, Software, Validation, Data curation, Visualization, Writing - review & editing, Supervision, Project administration.

## Supplementary information

### Pilot study overview

Prior to the experiment described in this paper we conducted a pilot study. Several parameters were different in the pilot. First, we used lower threshold estimates for (p_detected_ = 0.5 vs. 0.7 in the reported experiment). Second, the behavioral sampling frequency was higher, with 175 time bins 0.01 s (vs. 25 time bins of 0.05 s in the reported experiment). In the data of this pilot experiment we did not observe a significant increase in detection performance shortly after cue onset, i.e. a lack of the well documented cueing effect. We reasoned that the lack of this benchmark behavioral modulation could probably be attributed to a decrease in arousal and motivation of the participants for two main reasons. First, the combination of lower target luminance (p_detected_ = 0.5) and the same percentage of catch trials (i.e. 20%) resulted in (0.5*80% =) 40% of trials in which we expected participants to detect a target, i.e. in less than half of all trials. Second, due to the high amount of time bins the length of the pilot experiment was 4+ hours per participant (spread over two days). Because of this length, combined with the lower target luminance intensity, participants detected only few targets for a large amount of time. Hence, it is plausible that fatigue and lack of motivation affected their behavioral results (Morad, Lemberg, Yofe, & Dagan, 2000). Therefore, we decided to make changes to the experimental set up which led to the current experiment reported in the paper. However, to give a little more insight into the results of the pilot experiment, we summarize the methods and results here.

## Methods

There were 12 participants (main Experiment N = 20), of which one participant was excluded based on a low average performance (D’ = 0.71 for targets on a bright background and D’ = 0.41 for targets on a dark background). For the remaining participants detection performance was D’ = 2.13 (min. = 1.43, max. = 2.95) and D’ = 2.42 (min. = 1.16, max. = 3.21) for targets on a bright or dark background respectively. In the main experiment, these values were D’ = 2.47 and D’ = 2.66 on average for targets on a bright or dark background, respectively ([min, max] = [1.52, 3.26] and [1.65, 3.39], respectively). Of the remaining 11 participants in the pilot experiment, 4 were men (M = 21, SD = .71) and 7 were women (mean age = 25, SD = 9.83). All participants reported normal or corrected-to-normal vision.

### Difference with main experiment

Targets were presented in 175 CTOA’s in a window of 1.75 s after cue onset with intervals of 0.01 s (cf. 25 CTOA’s with 0.05 s intervals in the main experiment). Therefore, participants completed 1340 trials over 67 blocks which they could execute in two days. The threshold level in the staircase procedure was p_detected_ = 0.5 (cf. p_detected_ = 0.7 in the main experiment). Furthermore, the bright side of the screen was on the left side for all participants. For the analysis we used a temporal radius of 2 samples or 0.02 s (cf. temporal radius of 0.05 s in the main experiment). For the interaction between pupil and detection performance using a logistic linear mixed effects model per time point we performed 1000 permutations instead of 5000. All other aspects of the methods and analyses of the pilot study were identical to the main Experiment.

**Figure S1.**
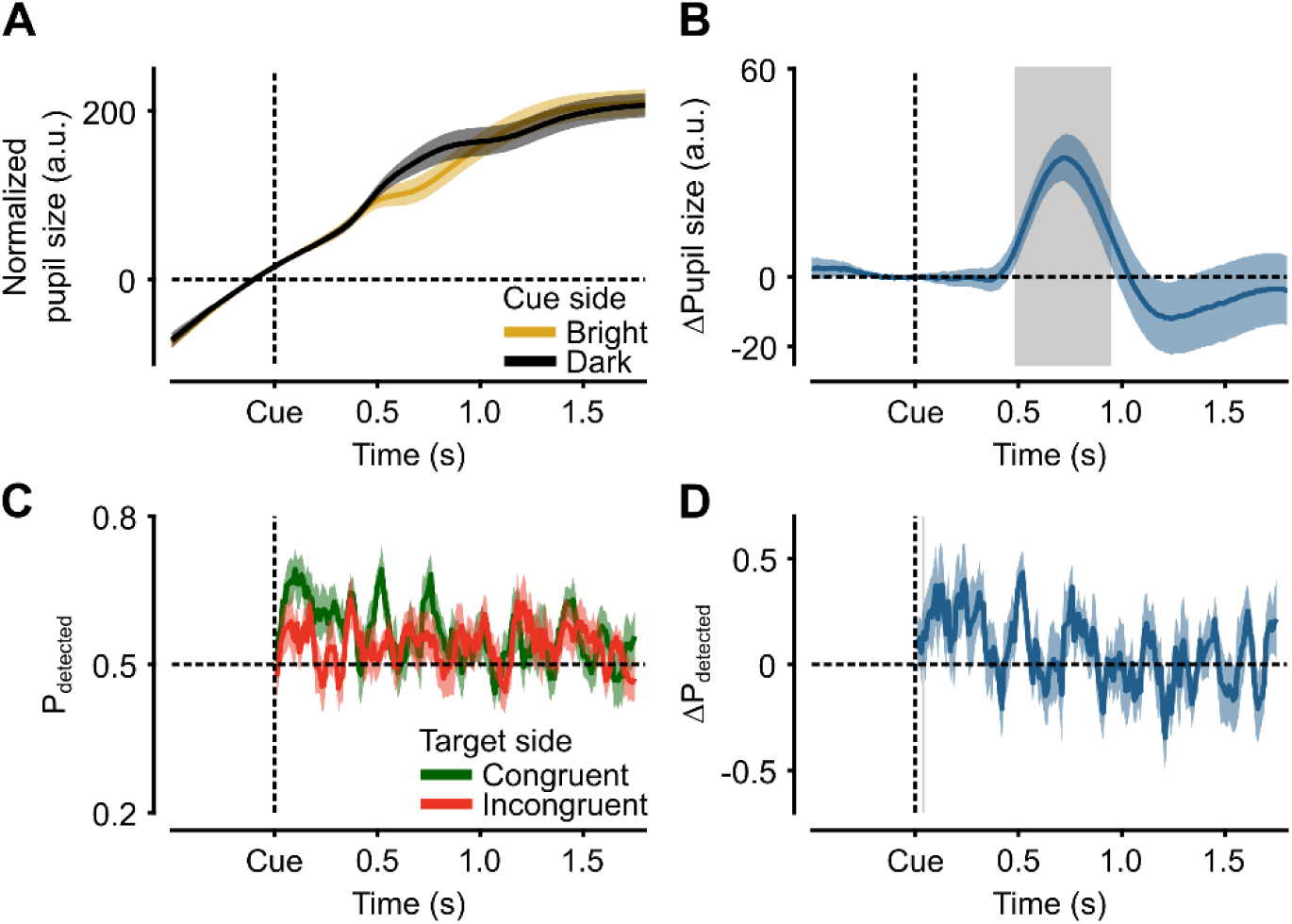
Results of the pilot experiment. **A**. Normalized pupil size (arbitrary units) as a function of time after cue onset (seconds) for trials in which the cue appeared at the bright (yellow) and dark (black) part of the screen. Shaded areas indicate standard error of the mean across participants. **B**. Relative pupil size (dark - bright). Grey patch (t = 0.49 - 0.94) indicates a period with a significant difference between pupil size dark and bright. **C**. Detection performance (proportion detected) for all 175 CTOA’s for congruent (green) and incongruent (red) trials. **D**. Cueing effect for detection performance (difference in proportion detected). Grey patch (t = 0.04) show a significant difference between congruent and incongruent trials.

### Pupillary cueing effect

In general, the pupil dilated before and after cue onset (Figure S1A). There was a significant modulation of pupil size by cue side between 0.49 and 0.94 s (maximum average = 34.4 a.u., p_cluster_ < 0.0001). After cue onset the pupil dilated more in the cue on dark condition compared to the cue on bright condition (Figure S1B).

### Behavioral cueing effect

For the detection performance of targets (Figure S1C), there was a significant positive cueing effects in favor of congruent trials only briefly after cue onset (Figure S1D, CTOA = 0.04, ΔP_detected_ = 0.10, p_cluster_ = 0.0062).

**Figure S2.**
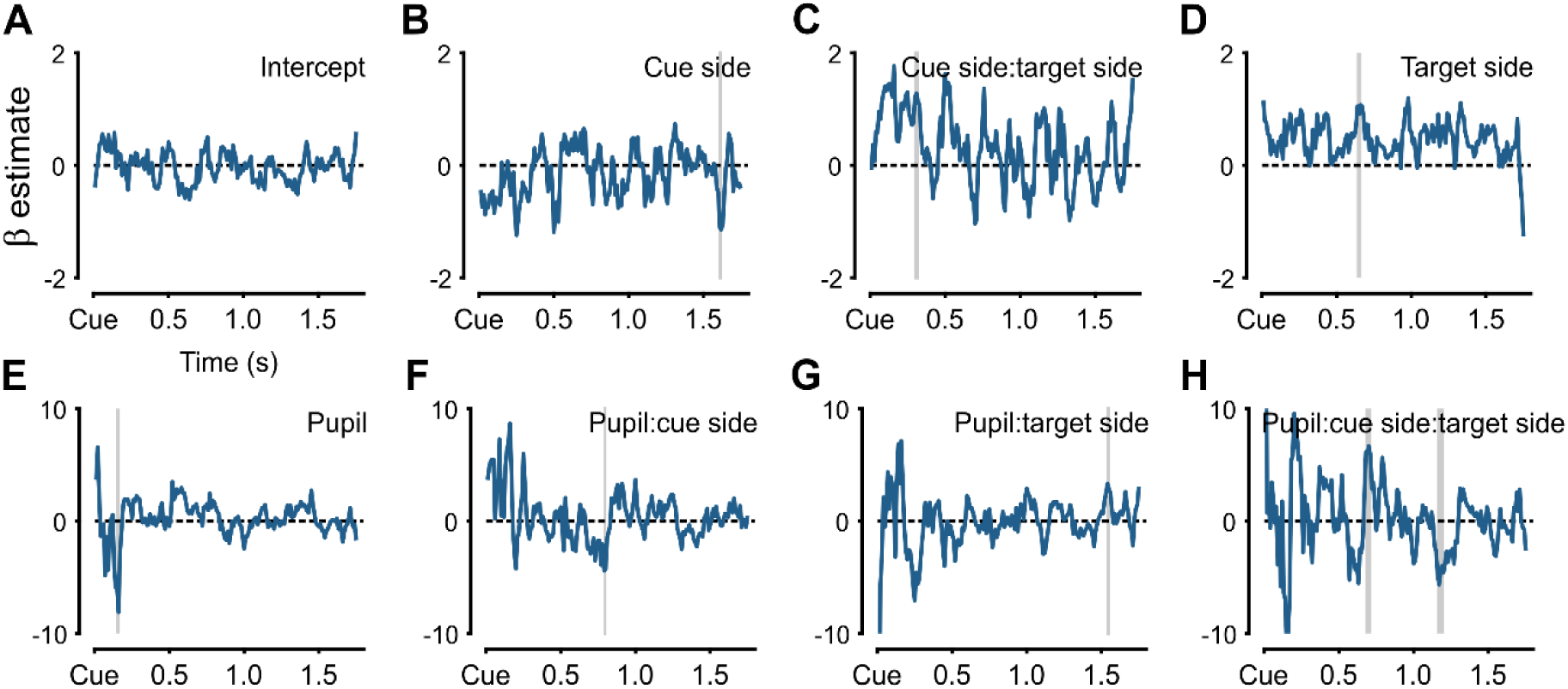
Result of logistic linear mixed effects model, modelling detection as a function of cue side (bright/dark), target side (bright/dark) and pupil size (a.u.). Models were made for each time point with a moving window of 5 time points (i.e. including the two preceding and two succeeding time points). Parameter estimates (β) are in log-odds relative to the intercept. Significance (vertical grey patches/lines) of the parameter estimates were corrected for multiple comparisons using threshold- free cluster-based permutation testing (N = 5000). **A**. Intercept, no significant deviations from chance level. **B**. Modulation of cue side. Significant effect at 1.61 s after cue onset. **C**. Modulation of the interaction between cue and target side. Significant effect from 0.30 to 0.32 s after cue onset. **D**. Modulation of target side. Significant effect from 0.64 to 0.66 s after cue onset. **E**. Modulation of pupil size. Significant effect at 0.15 s after cue onset. **F**. Modulation of the interaction between pupil size and cue side. Significant effect at 0.79 s after cue onset **G**. Modulation of the interaction between pupil size and target side. Significant effect at 1.54 s after cue onset. **H**. Modulation of the interaction between pupil size, cue side and target side. Significant effect from 0.68 to 0.71 s and 1.16 to 1.20 s after cue onset.

### Interaction between pupil size and detection

All the effects reported here are only significant at alpha = 0.05 for short durations and should therefore be interpreted carefully.

First, we re-analyzed the effect of target and cue side on detection performance using the Cue Side and Target Side variables in the mixed-effects model. The intercept (Fig. S2A), i.e. log odds of detection in trials where both the target and cue were presented on the bright side of the screen was never significantly different from chance level in the entire 1.75 s interval. Detection performance was significantly lower than the intercept at 1.61 seconds after cue onset when the cue was presented on the dark side and the target on the bright side (Fig. S2B). Detection performance was significantly higher than the intercept from 0.30 to 0.32 s after cue onset when both the target and cue were presented on the dark side (Fig.S2C). Detection performance was significantly higher than the intercept from 0.64 to 0.66 s after cue onset when the cue was on the bright side and the target on the dark side (Fig. S2D).

Next, we analyzed possible modulations of pupil size. Detection performance was lower at 0.15 s after cue onset with larger pupil size (Fig. S2E). Detection performance decreased with larger pupil size at 0.79 s after cue onset (Fig. S2F). Detection performance was increased with larger pupil size when the target was on the dark side, after a cue on the bright side at 1.54 s after cue onset (Fig. S2G). Detection performance was significantly increased with increased pupil size in trials where both the cue and target were presented on the dark side between 0.68 and 0.71 s after cue onset (Fig. S2H). The effect was reversed from 1.16 to 1.20 s after cue onset.

